# Generalized linear models provide a measure of virulence for specific mutations in SARS-CoV-2 strains

**DOI:** 10.1101/2020.08.17.253484

**Authors:** Anastasis Oulas, Maria Zanti, Marios Tomazou, Margarita Zachariou, George Minadakis, Marilena M Bourdakou, Pavlos Pavlidis, George M. Spyrou

## Abstract

This study aims to highlight SARS-COV-2 mutations which are associated with increased or decreased viral virulence. We utilize, genetic data from all strains available from GISAID and countries’ regional information such as deaths and cases per million as well as covid-19-related public health austerity measure response times. Initial indications of selective advantage of specific mutations can be obtained from calculating their frequencies across viral strains. By applying modelling approaches, we provide additional information that is not evident from standard statistics or mutation frequencies alone. We therefore, propose a more precise way of selecting informative mutations. We highlight two interesting mutations found in genes N (P13L) and ORF3a (Q57H). The former appears to be significantly associated with decreased deaths and cases per million according to our models, while the latter shows an opposing association with decreased deaths and increased cases per million. Moreover, protein structure prediction tools show that the mutations infer conformational changes to the protein that significantly alter its structure when compared to the reference protein.

## Introduction

First cases of the Severe Acute Respiratory Syndrome Coronavirus 2 (SARS-CoV-2) were reported from Wuhan, Hubei Province, China[1] last December 2019. Since then, SARS-CoV-2 has become the cause of the severe respiratory corona virus disease (COVID-19)[2], an epidemic initially declared in China[3], that over the course of 2-3 months spread globally to become a pandemic with grave repercussions (March 11th 2020, WHO publicly declared pandemic). The World Health Organization (WHO) reported: over 4.8 million confirmed cases of COVID-19, and over 316,000 deaths (report date: May 18th).

SARS-CoV-2 is an enveloped RNA virus, with a non-segmented, positive-sense (+ssRNA) genome of ~30 kb, amongst the largest identified *riboviria* RNA genomes[4,5]. It is believed to have originated from the bat coronavius Bat CoV RaTG13 (~96% identity)[6]. This is third outbreak of pathogenic zoonotic betacoronaviruses[6–10] within the last two decades, following the Severe Acute Respiratory Syndrome Coronavirus (SARS-CoV) in 2002[11], and the Middle East Respiratory Syndrome, MERS-CoV in 2012. However, these two predecessors were confined to epidemic proportions. So, what makes SARS-CoV-2 the cause of a pandemic unmatched by the previous two recent respirator syndromes (SARS-CoV and MERS-CoV)? The efficiency of SARS-CoV-2 can be attributed to a delicate balance between its transmission rate (high); and mortality rate (moderately high – compared to other two recent syndromes). It is difficult to accurately estimate differences in transmissibility and mortality because they are dependent on multiple factors which, vary significantly from one region to another. These factors include regional factors such as (i) access to testing, (ii) differences in clinical care, (iii) population demographic differences such as age and (iv) differences in weather conditions (e.g. heat and humidity). Nevertheless, current estimates for the basic transmissibility, commonly range between 2-5[12,13]. Estimates of mortality, deaths per confirmed cases average at 3.6% globally[14,15] SARS-CoV and MERS-CoV, although both had higher death rates than SARS-CoV-2 11%[16] and 35% mortality respectively[10,11], their transmission rates were significantly lower 1.5-2[12,17] and <1[17] respectively.

The transmission and mortality rates described above, provide clues as to how to study SARS-CoV-2 and devise strategies to investigate its genome taking under reconsideration its overall efficiency. Although the observed sequence diversity among SARS-CoV-2 strains is low, its rapid transmission rate provides ample opportunity for natural selection to act upon a range of viral mutations. High mutation frequency of a given position can be the consequence of one of two reasons: (i) either the mutation is in a neutral genomic region that allows for leniency in the frequency of changes with little or no impact virulence. (ii) The mutation is beneficial and actually provides a selective advantage to a viral strain enhancing its efficiency and overall propagation.

It is known that an “efficient” virus does not result in death for the underlining host. Therefore, the overall high transmission and moderate mortality rates of SARS-CoV-2 are what make this virus pandemic material.

The first SARS-CoV-2 strain to be sequenced was the index strain from Wuhan[18]. Currently the Global Initiative on Sharing All Influenza Data (GISAID) database, stores >23,000 SARS-CoV-2 strains (some of which are full-length sequences with high-coverage).

In this study, we capitalize on the ample amount of COVID information currently available through GISAID database and utilize it to develop a combined phylogenetic and statistical analysis that entails modelling the virulence of all mutations found across the SARS-CoV-2 strains sequenced to date. Our approach makes use of multiple sequence alignment (msa) tools (MAFFT)[19] and phylogenetic tree reconstruction methods (RAxML)[20] to obtain a map of all mutations in a variant calling format (vcf). The latter was done using snp_sites[21] tool maintained by sanger-pathogens, which characterizes genotypes for all the mutations (SNPs) found across all available sequenced strains in the msa. Our main objective is to highlight mutations which are associated with increased or decreased virulence, by using genetic data and countries’ regional information on deaths and cases per million. Initial indications of selective advantage of specific mutations can be obtained from calculating their frequencies across viral strains, which are obtained throughout the months of the COVID pandemic. These indicators include: (i) an increased frequency of strains that display a particular mutation over time in one region. (ii) Frequent recurrence of a specific mutation across different geographical regions. This would also be observed as nodes found on different branches of a phylogenetic tree. (iii) The existence of silent mutations, where a change at the nucleotide level (or codon), does not results in a subsequent change in the amino acid or the function of the overall protein. The threshold for identifying significant mutations can be set arbitrarily as done in[22] for the spike protein. However, by employing generalized linear models (GLMs) we reach an informed way to select mutations and provide additional information that is not evident from standard statistics or mutation frequencies alone. Subsequently, we select informative mutations and perform structural and conformational predictions of mutated and reference (according to initial Wuhan strain) proteins in order to investigate their effect on tertiary and quaternary protein structure. We propose two interesting mutations found in the nucleocaspid (N) (P13L) and ORF3a (Q57H) genes; that appear to be significantly associated with either increased or decreased deaths and cases per million according to our GLMs. Moreover, we show that the predicted conformations of the mutated protein are substantially different to the predicted reference structure.

## Results

### Statistical modelling for predicting key mutations for SARV-COV-2

The approach employed in this study provides an informed way of selecting mutations by applying GLMs fitted on various outcome and predictor variables. These include outcome variables: (i) deaths and (ii) cases per million information derived from the GISAID metadata file and predictor variables: (i) percent of occurrence of each mutation in strains found across different countries, (ii) binary mutation information (“0” absence of mutation, “1” presence of mutation), and (iii) country austerity measure response time. Our models are specifically designed to provide statistical evidence for fitting outcome variables that characterize the efficiency of the viral strains (e.g. death rates, transmission rates) with predictor variables such as the specific mutation occurrence found in populations and genotype information across all available strains. We are aware that genetic factors such as mutations frequencies are not the only variables that govern country death rates and transmission rates. Therefore, in addition to aforementioned predictor variables, we also consider the individual countries’ austerity measure response time as reported by the Oxford COVID-19 Government Response tracker data[23]. Two different types of models were constructed using: a) cases per million as the outcome variable and b) deaths per million as the outcome variable. Both generated unique results and ranked mutations in different order. Differences between model predictions provide valuable information on specific mutations (see next sections). We report the results from using both models types (a,b). Observing the top scoring mutations (see Tables 1 and 2) and analysing them with respect to their frequency across populations, provides valuable information with respect to their effect on viral virulence (see next section). In addition, excluding the response time from our predictor variables significantly alters the top ranked mutations and furthermore results in omission of some mutations from the top ranked significance lists seen in Tables 1 and 2 (see supplementary Table S1 and S2 for full list of ranked mutations using GLMs excluding response time from the predictor variables). To provide GLMs that characterize the circumstances responsible for changes in deaths and cases per million as accurately as possible, we decided to include response time as a predictor variable in all of our GLM fitting applications.

**Table 1.** Significant mutations according to our GLM of type a, which utilizes cases per million as outcome variable and including response time as a predictor variable. Two protein mutations of interest are and highlighted in light blue (see sections below for details).

| Mutation ID | Codon | Genomic Position | Gene | P-value |
| --- | --- | --- | --- | --- |
| Mut9699 | 13 | 28311 | N | 0.002194766 |
| Mut8612 | 57 | 25563 | ORF3a | 0.007292035 |
| Mut7944 | 614 | 23403 | S | 0.007319234 |
| Mut1515 | 924 | 3037 | ORF1ab | 0.007523798 |
| Mut5088 | 314 | 14408 | ORF1ab | 0.00818189 |
| Mut766 | 265 | 1059 | ORF1ab | 0.008613815 |
| Mut4923 | 88 | 13730 | ORF1ab | 0.013379563 |
| Mut8101 | 789 | 23929 | S | 0.016250144 |
| Mut2610 | 2016 | 6312 | ORF1ab | 0.017059216 |

**Table 2.** Significant mutations according to our GLM of type b, which utilizes deaths per million as outcome variable and including response time as a predictor variable. Two selected protein mutation of interest highlighted in light blue (see sections below for details).

| Mutation ID | Codon | Genomic Position | Gene | P-value |
| --- | --- | --- | --- | --- |
| Mut1515 | 924 | 3037 | ORF1ab | 7.26E-05 |
| Mut7944 | 614 | 23403 | S | 7.71E-05 |
| Mut5088 | 314 | 14408 | ORF1ab | 0.000115 |
| Mut3461 | 2839 | 8782 | ORF1ab | 0.003728 |
| Mut9636 | 84 | 28144 | ORF8 | 0.003881 |
| Mut4154 | 3606 | 11083 | ORF1ab | 0.004302 |
| Mut9699 | 13 | 28311 | N | 0.00463 |
| Mut8877 | 251 | 26144 | ORF3a | 0.006989 |
| Mut5205 | 446 | 14805 | ORF1ab | 0.008682 |
| Mut5329 | 619 | 15324 | ORF1ab | 0.019863 |
| Mut8101 | 789 | 23929 | S | 0.021618 |
| Mut4923 | 88 | 13730 | ORF1ab | 0.022187 |
| Mut2610 | 2016 | 6312 | ORF1ab | 0.022238 |
| Mut9924 | 203 | 28881 | N | 0.035819 |
| Mut9925 | 203 | 28882 | N | 0.036078 |
| Mut9926 | 204 | 28883 | N | 0.03638 |
| ... | ... | ... | ... | ... |
| Mut8612 | 57 | 25563 | ORF3a | 0.479085 |

### Analysis of model results

The top ranking mutations highlighted by our GLMs that have been the centre of multiple recent research studies[22,24,25]. These include mutations that appear as major high frequency peeks in genomic plots for the SARS-CoV-2 virus strains[26]. Notable examples include: 1) Mutations at position 14408 (codon 314), 3037 (codon 924) and 11083 (codon 3606) on the ORF1ab[25], that show a high frequency of occurrence (0.65, 0.65 and 1.3 respectively) across viral strains sequenced to date. 2) The very cosmopolitan mutation on position 23403 (codon 614[22], frequency 0.65) of the S protein (see Fig 1). 3) The highest frequency peeks found in the N protein at positions 28881-28883[25] (codons 203, 204 respectively). 4) Well known high frequency mutations found in the ORF3a gene such as mutation at position 26144 (codon 251)[25].

**Fig 1.**
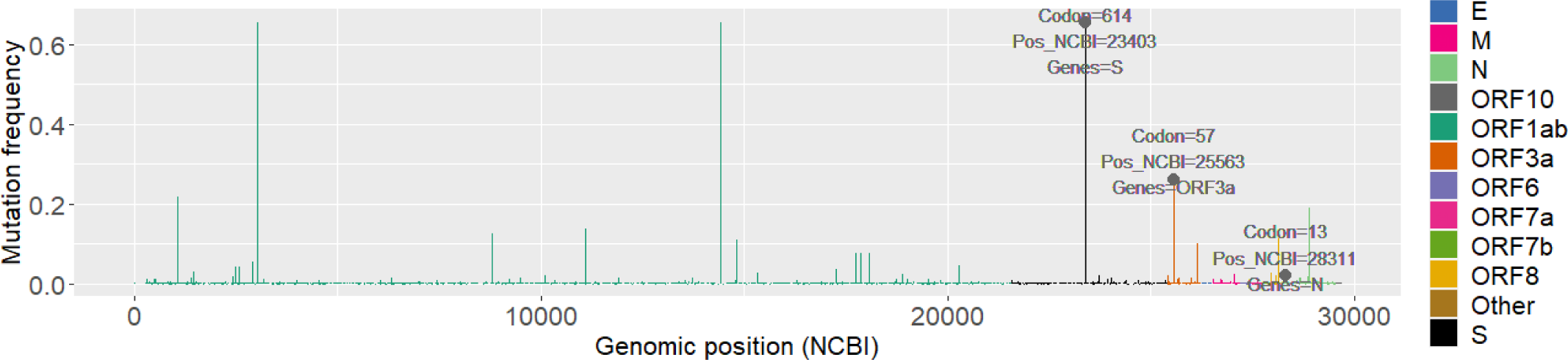
Global genomic frequency plot. Mutations of interest, P13L and Q57H, are labelled in the plot together with the D614G mutation for comparison.

### Mutations of interest for studying the efficiency of SARS-COV-2

What we consider as the most significant contribution of our model, is that it also highlights less studied mutations that appear to correlate with death and contamination rates. We highlight two mutations of relatively low global frequencies (see Fig 1): 1) Mutation at position 28311 characterized by a P to L change on codon 13 (P13L) in the N protein. 2) Mutation at position 25563 on codon 57 of the ORF3a protein characterised by a Q to H (Q57H) amino acid change. These mutations were selected because of their highly significant ranking according to our GLMs (see Tables 1 and 2). In the case of P13L, this mutations attained high significance (*p-values*: 0.002 and 0.005) according to our models. In addition, what rendered this mutation interesting for further analysis, was its negative correlation with both deaths and cases per million (see Fig 2A,B). Albeit it showed a lower frequency of occurrence across different countries and in the general population (see Fig 1). In the case of Q57H, this mutation showed high significance according to type a models (*p-value*: 0.007) however, it was not deemed significant according to type b models. It was, however, selected due to its unique correlation patterns showing opposing decreasing and increasing death and transmission rates respectively (see Fig 3A,B). Q57H has an elevated frequency level across countries and in the overall population (see Fig 1). Our highlighted mutations were also compared to the well-studied D614G mutation in our consecutive detailed analyses (see Fig S1).

**Fig 2.**
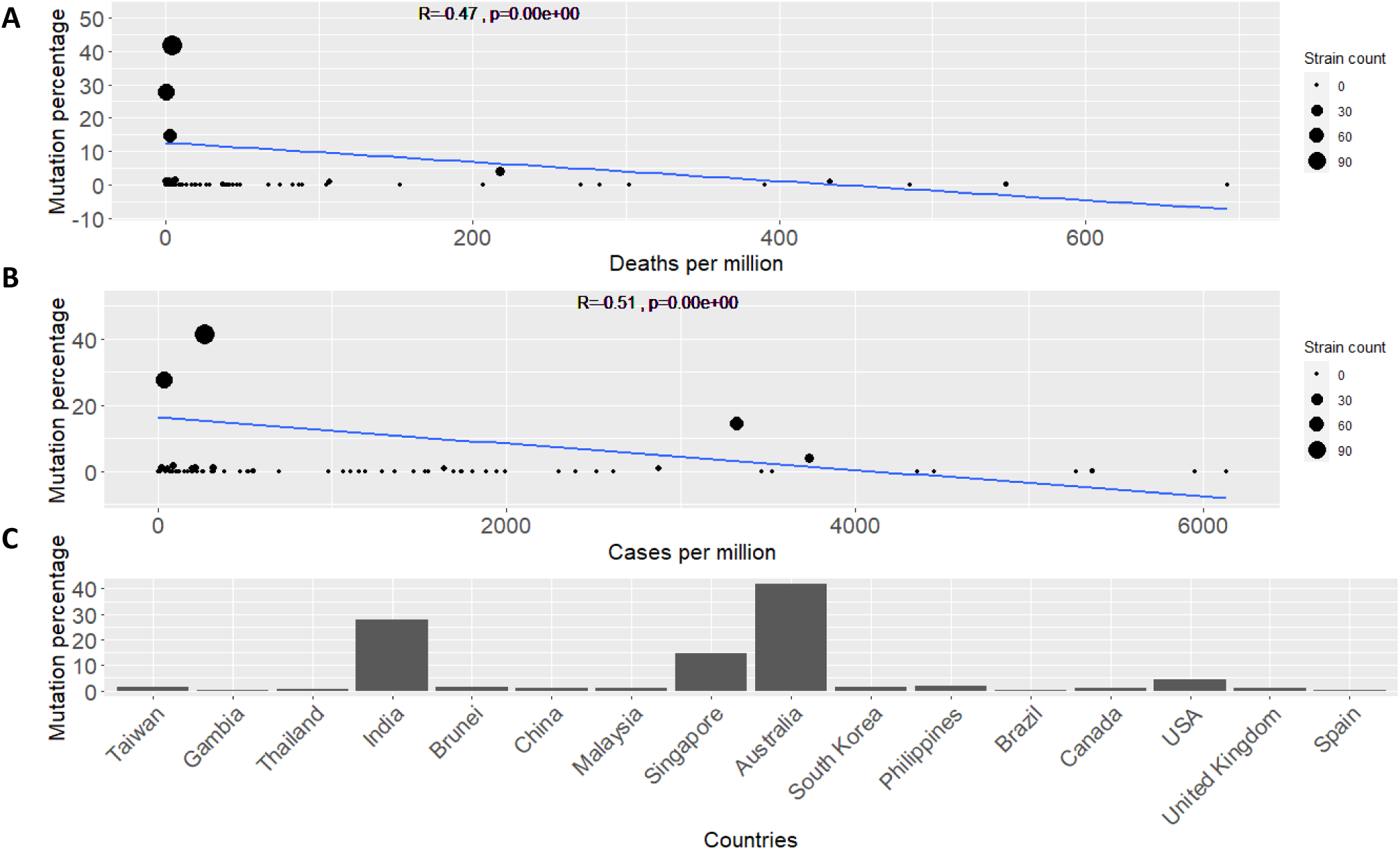
Analyses plots for N protein mutation at position 28311 (P13L) **A**. Regression model line showing the simplified fit for mutations percentage across countries and the deaths per million for each country. Pearson’s correlation is shown by the *R* value accompanied by the *p*-value of the correlation coefficient. **B**. Similar regression fit for mutations percentage across countries this time showing cases per million for each country. Pearson’s correlation is shown by the *R* value accompanied by the *p*-value of the correlation coefficient. **C**. Detailed histogram of the percentage occurrence of the mutation across different countries. These values make up the “percent” predictor variable utilized in our model (see Materials and Methods). Countries are sorted with increasing deaths per million.

**Fig 3.**
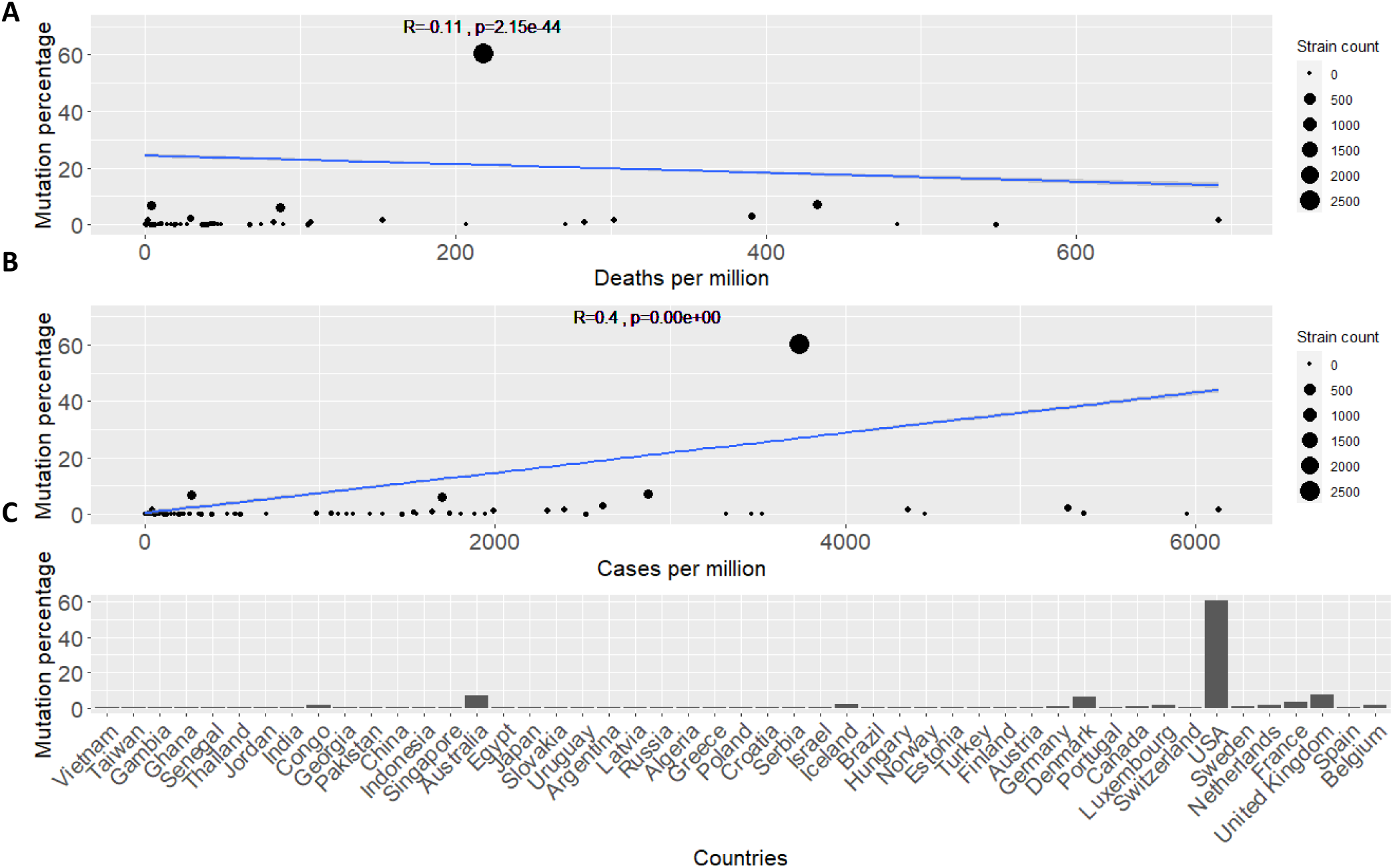
Analyses plots for ORF3a protein mutation at position 25563 (Q57H) **A**. Regression model line showing the simplified fit for mutations percentage across countries and the death rate per million for each country. Pearson’s correlation is shown by the *R* value accompanied by the *p*-value of the correlation coefficient. **B**. Similar regression fit for mutations percentage across countries this time showing cases per million for each country. **C**. Detailed histogram of the percentage occurrence of the mutation across different countries. Countries are sorted with increasing deaths per million.

### The P13L mutation tracking analysis

The P13L mutation is characterized by a change from C-to-T,Y(C or T),G base change at position 28311 in the Wuhan reference strain. It is a highly significant mutation according to our GLMs and moreover appears to be significant even when taking in consideration the country austerity measures response time. P13L is being tracked by the GISAID, we refer to the clade of sequences that constitute this mutation as the “P13L” clade (see Fig S2 for the P13L phylogenetic clade position on our ML tree). It is present in 1.9% of the global samples, and shows a global distribution appearing in Asia, America and Australia and the UK. It has not yet been analysed in depth and we propose an important role for this mutation in viral stability and overall virulence. The data available for study (12/05/2020), are in accordance with a founder effect in Asia, followed by an increase in Australian samples. During March and April, sampling of the data from GISAID showed that the mutation frequency was further spreading in Asia and then shifting to the UK and the Americas. The observed geographic spread is consistent with that of a mutation showing early signs of an expanding global frequency (Fig 4A).

**Fig 4.**
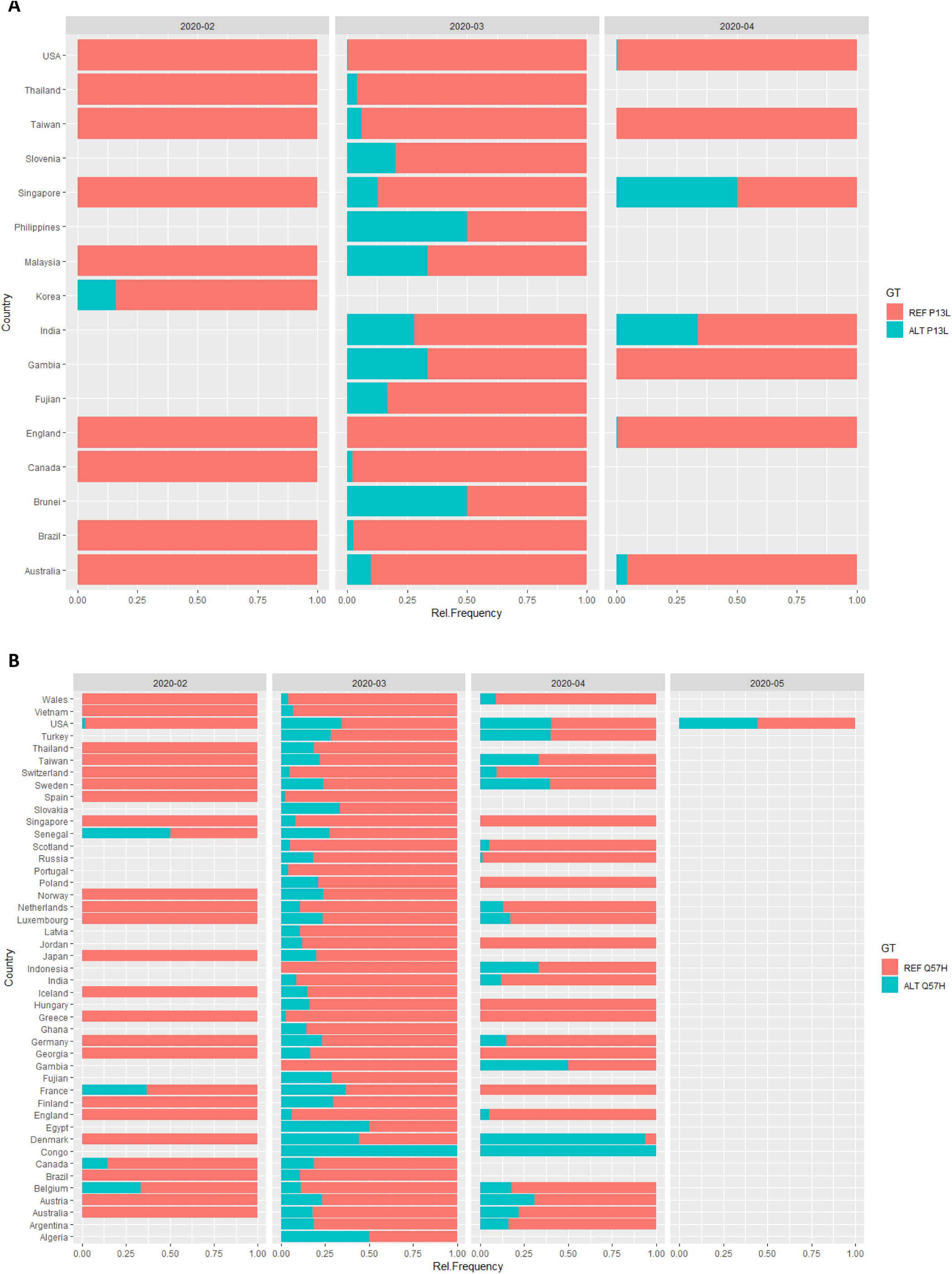
Mutation tracking relative frequency plots. **A**. P13L transmission in months from its first occurrence in Asia (Korea) to its transmission across other countries and continents. **B**. Q57H transmission in months from its first occurrence in Europe (France) to its transmission across the globe.

Mutation tracking across countries with available adequate sampling, revealed a consistent pattern of mutation spreading. In countries where sequences were sampled before March (e.g. Korea, Australia, UK, USA), the reference P13L (REF-P13L) form appeared to be dominate in this early period of the pandemic (Fig 4A). The introduction of alternate P13L (ALT-P13L) in the population, was followed by a rise in its frequency, and in some cases ALT-P13L constituted ~50% of the population (Fig 4A).

In the UK, the ALT-P13L form began circulating later in the pandemic (April). Through March, ALT-P13L became documented in South America, and it constituted a substantial percent of contemporary sampling (Fig 4A). In North America, initial infected samples across the continent showed an establishment of the original REF-P13L form, as the pandemic progressed, the ALT-P13L appeared in both USA and Canada, becoming a well-documented form in both nations by the end of March.

Australia follows a similar transition pattern, from REF-P13L to ALT-P13L, as described above, however the introduction of the mutation in the Australian population appears to have preceded that of USA and Canada. In fact the earliest ALT-P13L mutation outside of Asia was identified in Australia (NSW25.2020.EPI_ISL_417388.2020.03.05.Oceania). This can account for the much higher frequencies (~10 times) of the ALT-P13L form observed in Australia compared to the former two countries. Most Asian samples showed a dominance of the original Wuhan REF-P13L form in early in the pandemic (February), however for Asian countries outside of China, the ALT-P13L form was well-documented and becoming established by March (Fig 4A). It is difficult to track the P13L mutation in China due to the lack of available Chinese sequences in GISAID after March 1st. Sparse sampling, as observed in South America and Africa, can affect mutation tracking, and this has to be taken under consideration when analysing frequency plots (Fig 4A). In summary, our analysis of the data shows that the REF-P13L form may be under selective advantage as shown by persistent, recurring transitions to the ALT-P13L form in multiple regional geographical location throughout the pandemic.

### The Q57H mutation tracking analysis

The Q57H mutation has previously been studied in detail[24] so we do not perform a detailed mutation tracking for this mutation. For completion we show relative frequencies of the mutation across time (Fig 4B) consistent with its founder effect in Europe and its transmission across the globe. Although previously studied, the Q57H mutation has not been associated with virulence and the precise mechanism for its selective advantage and increased frequency across populations has not been investigated. It has a relatively high frequency of occurrence according to the GISAID data analysed in this study (>0.26). The observed opposing correlations with deaths and cases reported per million according to demographic data, presented in our study may provide clues for the increased frequency of this mutation. As previously mentioned, the choice for highlighting this mutation is: 1) a consequence of the significant scores obtained by Q57H by both our model types (see Tables 1 and 2) its unique pattern of correlation with respect to death and transmission rates. Notably, this mutation is one of the few (4/100) top significant mutations that exhibit a negative correlation with deaths per million and a positive correlation with cases per million. This is a suitable pattern for delineating viral mutation fitness, as preferably a competent virus should have high transmission rates and relatively moderate death rates to secure its persistence.

For comparison, mutation tracking of the well-studied D614G mutation[22] is also performed (Fig S3).

### P13L and viral transmission and virulence

We hypothesize that if the P13L mutation can affect transmissibility and death rates, it might also impact severity of disease. High-throughput clinical outcome data for COVID are not readily available at the moment, therefore we focused on geographic regional data reporting death and transmission rates according to the Worldometers.info website. Populations harbouring the mutation show a higher mean death rate compared to the converse. Statistical analysis of the populations with and without the P13L mutations shows substantial evidence that the two distributions are significantly different (Wilcoxon test – *p*-value 2.2e-16) (see Fig 5A). This is in contrast to the results obtained by our model. This discrepancy can be attributed to the differences between performing univariate statistics to using GLMs. Our models have the additive advantage to incorporate multiple quantitative factors in consideration, such as the percent value for each country, while box plot statistics only looks at presence or absence of mutation in a population. For example Spain has a high death rate shifting the mean up but only a small percent (0.35%) of the mutation is actually present throughout the population. Our GLMs are specifically suited to quantify this type of information and provide a more accurate depiction of the quantitative realm studied here. For comparison reasons we also perform the analysis performed on the D614G mutations, which shows similar results, with the population comprising the mutation showing significantly different (Wilcoxon test – *p*-value 6.1e-15) increased death rates compared to the referenced-based population (see Fig S4). Similar phenomena where observed when comparing results using the cases per million parameter to assess the different distributions (see Fig 5B, and Fig S4B).

**Fig 5.**
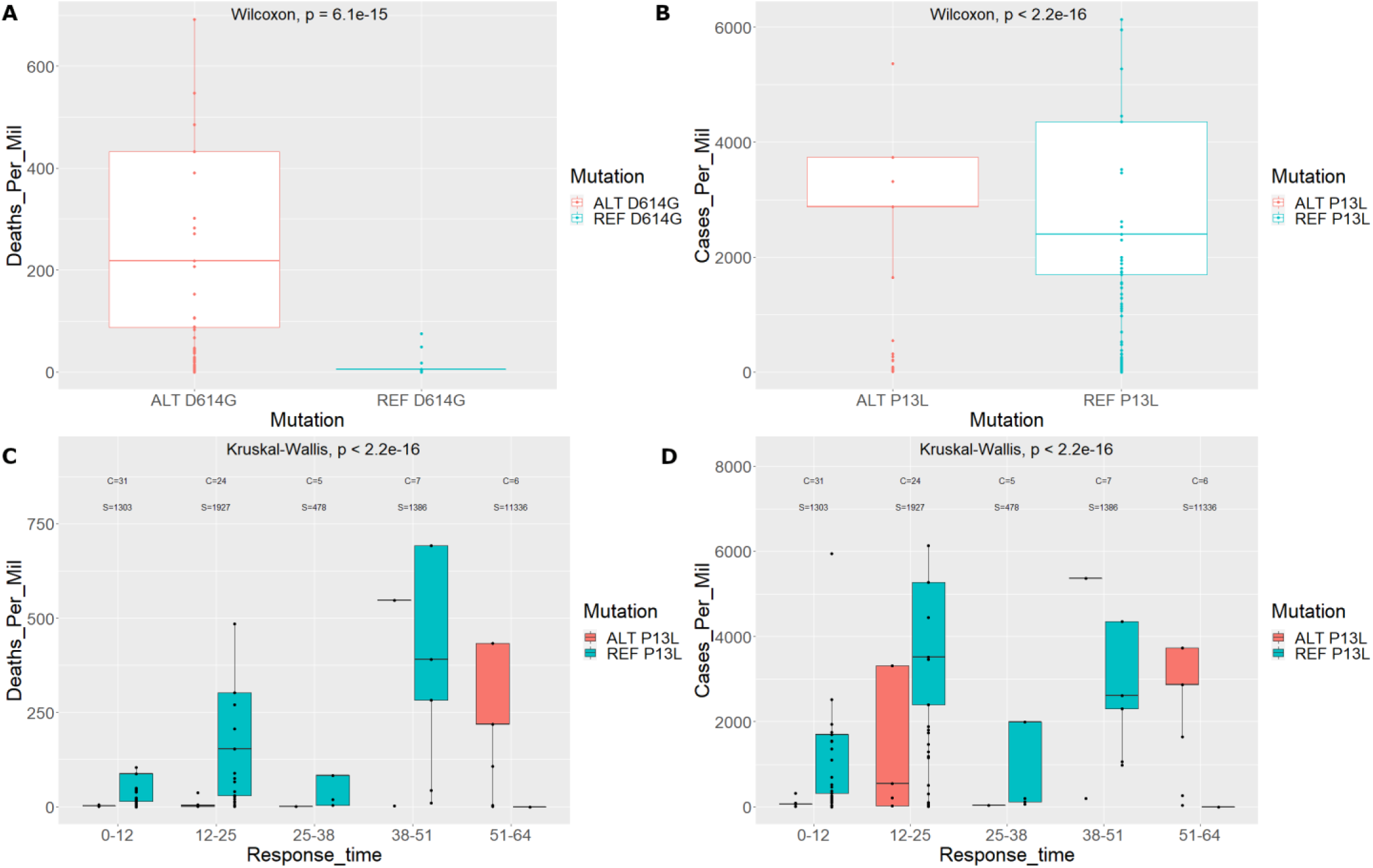
Boxplot distributions with and without the P13L mutation. **A.** Deaths per million for countries with the P13L mutation and the reference mutation. **B.** Cases per million for countries with the P13L mutation and the reference mutation. Keep in mind that each data point can represent more than one country. **C.** Deaths per million for countries with the P13L mutation and the reference mutation including response time separation. *C* denotes the number of unique countries in the group and *S* is the number of strains in the group. **D.** Cases per million for countries with the P13L mutation and the reference mutation including response time separation. *C* and *S* are as denoted for panel **C**.

### Q57H and viral transmission and virulence

The transmissibility and death rates patterns of Q57H mutation are consistent with the high contamination and low lethality of SARS-CoV-2. Geographic regional data reporting deaths and cases per million show that populations harbouring this mutation exhibit higher mean death rates compared to reference populations. However, statistical analysis of the deaths for populations with and without the Q57H mutations shows evidence that the two distributions are marginally not different (Wilcoxon test – *p*-value 0.069 - see Fig 6A). The converse was observed when comparing results using the cases per million parameter to assess the different distributions (Wilcoxon test – *p*-value 1.6e-8 - see Fig 6B). Results here are in total agreement to our model which predicts a correlation with mutation percent found in each country and cases per million, while a negative correlation with the deaths per million.

**Fig 6.**
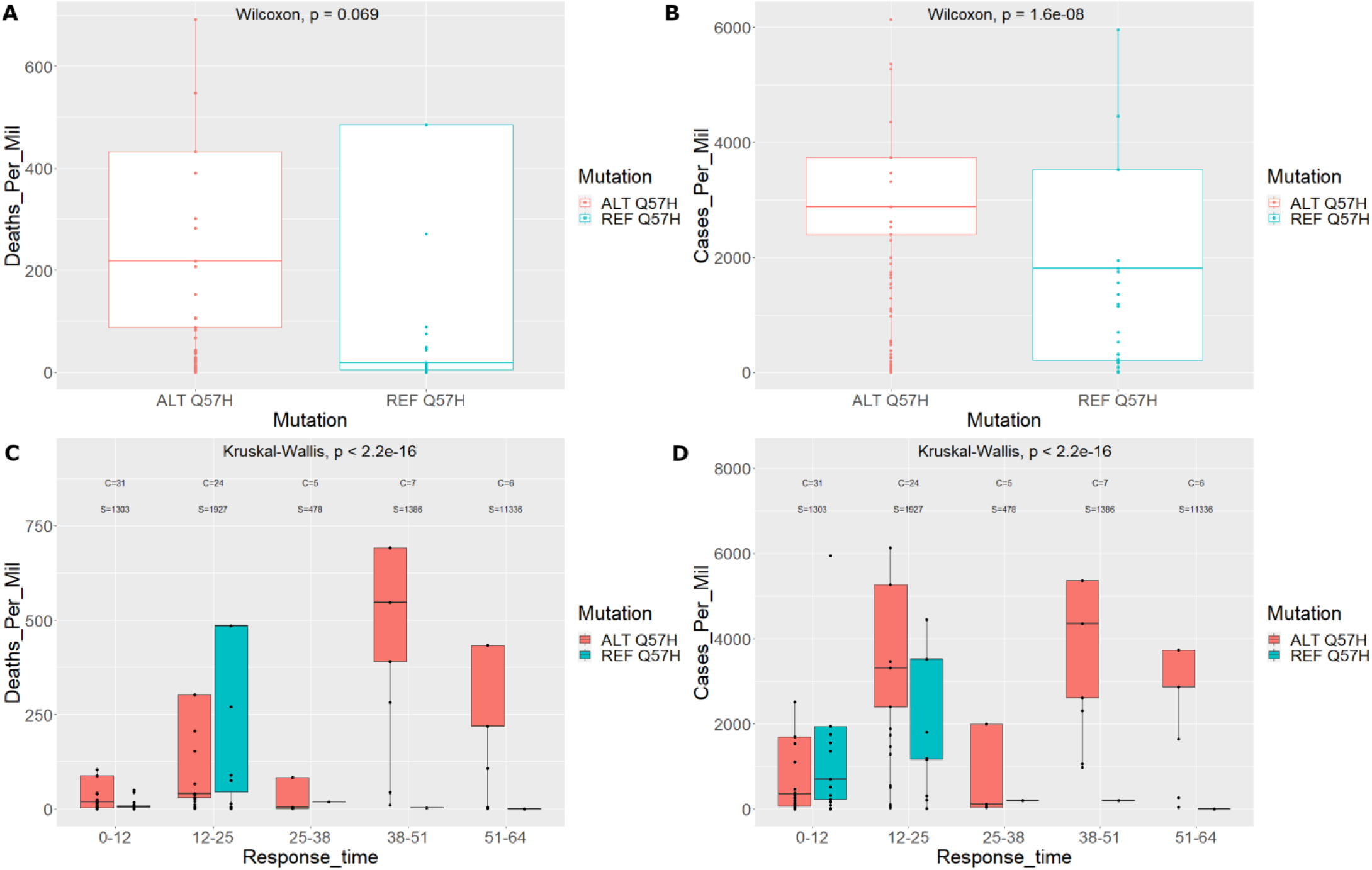
Boxplot distributions with and without the Q57H mutation. **A.** Deaths per million for countries with the Q57H mutation and the reference mutation. **B.** Cases per million for countries with the Q57H mutation and the reference mutation. **C.** Deaths per million for countries with the Q57H mutation and the reference mutation including response time separation. *C* denotes the number of unique countries in the group and *S* is the number of strains in the group. **D.** Cases per million for countries with the Q57H mutation and the reference mutation including response time separation. *C* and *S* are as denoted for panel **C**.

To investigate patterns of the effect of country response time with respect to the death and transmission rates as well as the mutation genotype, we performed a similar distribution analysis including the response time in months. The overall analysis for the Q57H mutation showed both death and transmission rates increased with response times, however there was significant drop in deaths per million for populations with the ALT-Q57H compared to cases per million (see Fig 6C,D). This decrease in death rates was especially documented for the 12-25 month response group, which also comprised of most of the countries studied (see Fig 6C). A phenomenon which was observed at a much lesser effect when investigating cases per million (see Fig 6D). Moreover, it is clear that countries that delayed their response time ultimately attained more cases and deaths per million[23]. It is evident that these countries also had a higher occurrence of the ALT-Q57H mutation. This can possibly be attributed to the increased number of cases in these countries, providing the virus with greater opportunity for natural selection to act upon this specific mutation.

### P13L and potential mechanisms for enhanced fitness

In order to provide an explanation for the reasons why the P13L mutation appears to be associated with decreased transmission and death rates, we turn to structural prediction and docking tools. P13L is located on the surface of the Nucleocapsid (N) protein known to form helical ribonucleocapsids (RNPs) with the positive-sense, single-stranded RNA genome of the SARS-CoV-2 virus. It also interacts with virus membrane (M) protein and subsequently plays a fundamental role during virion assembly. The QHD43423 structure of the N protein and QHD43419 structure of the M protein, were retrieved from the I-TASSER repository containing the 3D structural models of all proteins encoded by the genome of SARS-CoV-2. Structures of the REF-P13L and ALT-P13L mutations, exhibited significant differences in protein stability, with the later presenting a decrease in stability, compared to the REF-P13L compound (*ΔΔG* = 0.629±0.012kcal/mol). Both amino acids are exposed to the surface with RSA values 76.4% and 75.6% for the neutral Proline (P) and hydrophobic Leucine (L) residues, respectively.

The native 7mer-RNA duplex (PDB ID: 4U37) was used for RNA-protein docking, as described by Dinesh et al.[27]. RNA-protein docking with HDOCK and MPRDock algorithms, revealed changes on the assembly of RNP complexes (RMSD 2.719±3.068Å) (Fig 7A,B). Investigation of the N-M interaction was also carried out. The REF-P13L – M protein complex exhibited an increase of the binding affinity (ΔG -13.050±0.495kcal/mol and Kd 2.95E-10±2.192E-10M) compared to the ALT-P13L – M protein complex (ΔG - 12.750±0.212kcal/mol and Kd 4.600E-10±1.270E-10M). In addition, structural changes are evident as demonstrated upon structural alignment (RMSD 1.018±1.027Å) (Fig 7 C,D).

**Fig 7.**
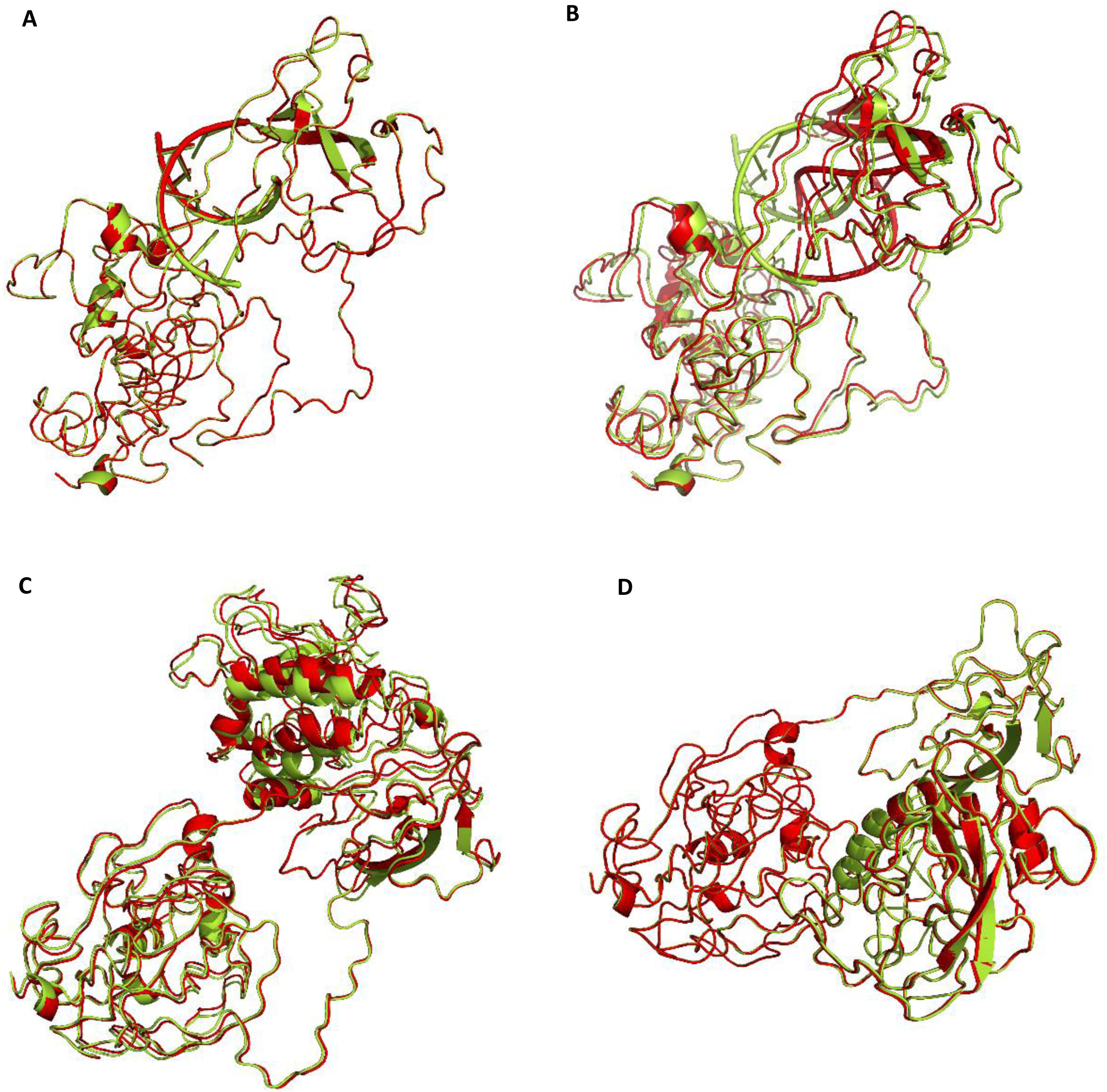
Aligned tertiary and quaternary structural predictions of the complex between the Nucleocapsid (N) protein and RNA (PDB ID: 4U37). **A**. Protein-RNA HDOCK docking simulations’ result. **B**. Protein-RNA MPRDock docking simulations’ results. REF-P13L complexes colored in red, ALT-P13L complexes colored in green. **C.** Protein-protein ClusPro docking simulations’ result of aligned tertiary and quaternary structural predictions of the complex between the Nucleocapsid (N) protein and membrane (M) proteins. **D.** Same analysis shown in panel **C** performed using protein-protein HDOCK docking simulations. REF-P13L complexes colored in red; ALT-P13L complexes colored in green.

### Q57H and potential mechanisms for enhanced fitness

Structural prediction analysis for the Q57H mutation provides evidence to support its association with decreased death rates and increased transmission. The ORF3a protein forms homotetrameric potassium sensitive ion channels (viroporin) and may modulate virus release. The Q57H mutation is located on the outer surface of the protein and specifically in the binding domain that is directly involved in the tetramerization process. We performed a structural analysis using I-Tasser[28] and created high quality tertiary and quaternary structure predictions for the ORF3a protein (C-score=-3.14, Estimated TM-score = 0.36±0.12, Estimated RMSD = 13.5±4.0Å). The Gln57 residue is a charge-neutral, polar amino acid, which resides 2 amino acids far the transmembrane region (35-55) of the ORF3a protein. The ALT-Q57H is defined by a substitution to a positively-charged and polar Histidine (His). Comparison of the structures obtained using the REF-Q57H mutation and the ALT-Q57H (see Fig 8A,B) show significant differences both in structural conformation as well as stability. With the later exhibiting a more stable (−29.0ΔG, Kd = 5.6E-22M) structure affinities for the overall ORF3a viroporin compared to the former (−26.8ΔG, Kd = 2.3E-20M). Further, in depth molecular analysis reveals disruption of hydrogen bonds and hampering and shifts in other inter-molecular interactions in the ALT-Q57H compared to the REF-Q57H structure of the protein (see Fig 8 C,D). The ORF3a protein is known to be involved in the pathway implicated in pore formation in the host cellular membrane and is crucial in viral budding and release into the external cellular space. The generation of a more stable structure that can ultimately lead to a more sustainable viral cycle can be the functional aetiology for observing a greater correlation with transmission of the virus. However, the fact that the ORF3a protein is not involved in host receptor recognition and binding, could be the reasoning behind the observation that overall death rates are not increased as a consequence of this mutation.

**Fig 8.**
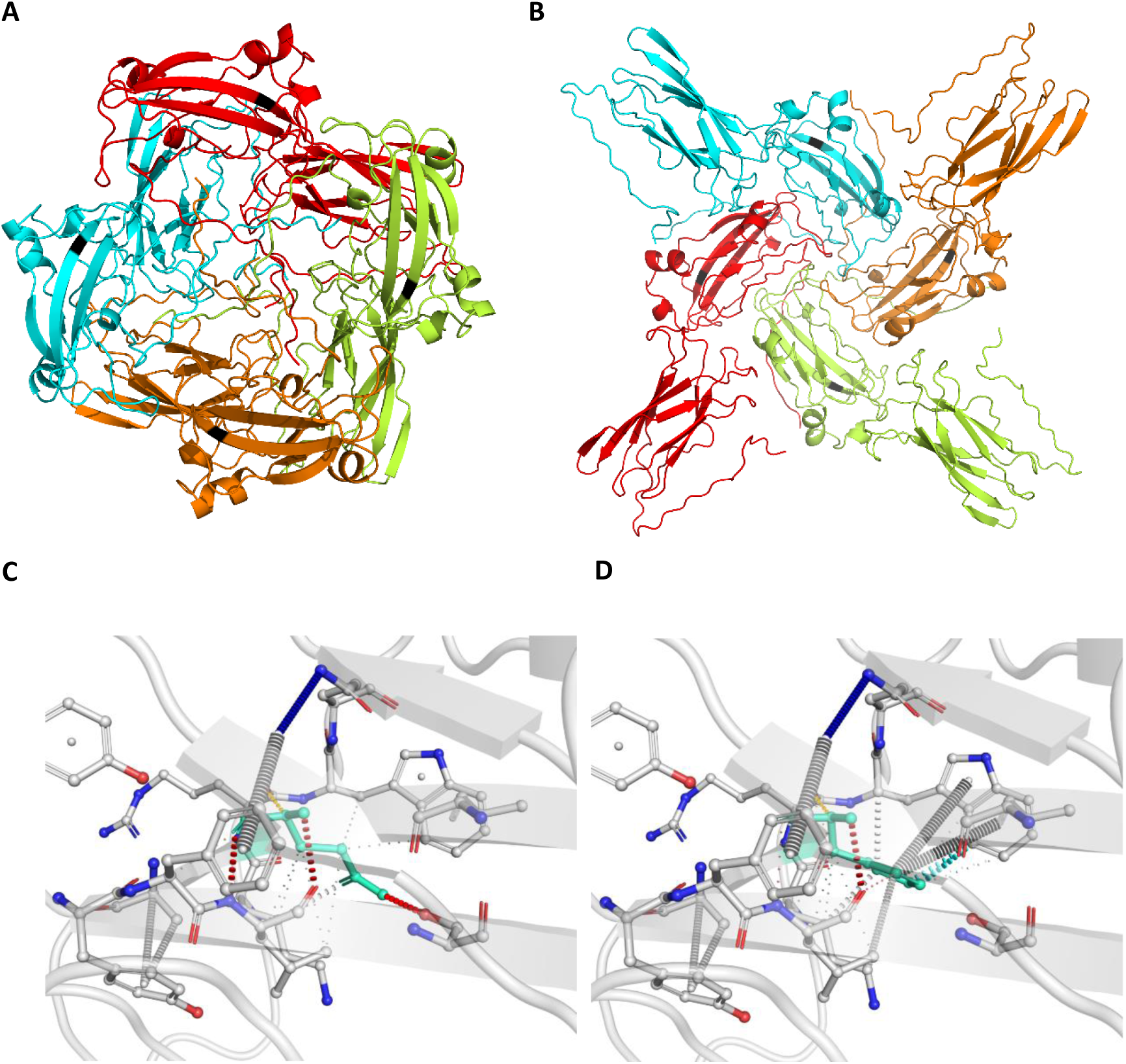
Tertiary and quaternary structural predictions of ORF3a protein. **A**. Structural prediction with REF-Q57H. **B**. Structural prediction with ALT-Q57H. The location of the 57th amino acid in both structures is denoted in grey. **C**. Detailed molecular structural conformational changes of ORF3a protein showing the REF-Q57H. **D**. Same molecular analysis for the structure with ALT-Q57H. Hydrogen bonds in red, weak hydrogen bonds in orange, halogen bonds in blue, ionic interactions in yellow, aromatic contacts in light-blue, hydrophobic contacts in green, carbonyl interactions in pink, VdW interactions in grey.

## Discussion

Investigating viral strain mutation virulence can be a delineating factor in predicting viral transmission and mortality at a population level. Diagnostic tests that can sequence targeted viral genomic regions for specific mutations can provide invaluable information to countries for taking fast and efficient actions in dealing with a pandemic such as that created by the SARS-CoV-2. This study shows that regression models can be used to provide information at the SNP level that allows inferences on how mutations are correlated with death rates and transmission rates. Frequency level information can only provide information on more cosmopolitan mutations and less frequent mutations are largely neglected. The use of generalized linear regression models presented in this study allows for the assessment of all mutations irrespective of their current frequency, highlighting mutations of interest that are correlated or anti-correlated with viral virulence. This permits inference on mutations that are on the rise and potentially can alter strain virulence affecting the overall progression of the virus across populations. Moreover, external (non-genetic factors) such as countries’ response times to reach stringent austerity measures can be incorporated in the models, allowing for an accurate examination of the genetic factors under investigation. A less frequent mutation such as the P13L mutation on the N protein of SARS-CoV-2, is highlighted using the GLMs constructed in this study and shows a negative correlation with respect to country deaths and cases per million. The crystal structure of the N protein has been the focus of other studies revealing potential drug targeting sites[29]. RNA-protein docking performed in this study show a potential impact of the ALT-P13L on RNA binding affinity. Furthermore, investigation of the N-M interaction reveals that the ALT-P13L – M protein complex exhibited a decrease in binding affinity. These observational allude to a possible affect on virion assembly.

The more frequent but not so well studied Q57H mutation shows patterns that are in accordance to optimal viral fitness and persistence. At present time statistical analysis comparing different populations, with and without the aforementioned mutation, appear to agree with our GLMs in accordance to non-parametric tests, it will be interesting to track this mutation in future population distributions and examine whether they will exhibit similar patterns. Moreover, structural analysis of the Q57H shows significant conformational changes in the ORF3a protein. According to free energy calculations, these are potentially accountable for a more stable quaternary structure of the protein, possibly leading to an enhanced release of viral particles, in accordance to its role in host cell pore formation and budding.

Taking the frequently occurring D614G as an example it is evident that this mutation is becoming the dominant mutation across populations and its rapid spread denotes a natural selection pressure that shows increased fitness for this mutation. Fitness for a viral mutation can be governed by factors such as increase of viral transmission, such as that observed for D614G. However, a successful virus will not evidently kill its host, so increased contamination rates have to be coupled with decreased deaths rates. This is exactly what is observed for the Q57H and P13L mutations, which provide a counter balance to mutations (e.g. D614G) that are positively correlated with both death rates and transmission rates.

## Methods

### Datasets

Sequence data for all available SARS-COV-2 strains were obtained from The Global Initiative for Sharing All Influenza Data (GISAID), at https://gisaid.org. Individual country austerity measure responses were obtained from the Oxford COVID-19 Government Response tracker data[23].

### Overall approach

Full genomic SARS-CoV-2 sequences of high sequencing resolution were obtained from GISAID. The nextstrain[26] pipeline was downloaded locally and the commands for filtering, aligning and constructing phylogeny were used according to the nextstrain’s best practices. MAFFT[19] was used to construct a multiple sequence alignment (MSA). Phylogeny was estimated using the RAxML[20] maximum likelihood algorithm for phylogenetic tree construction. The vcf was generated using the snp_sites tool available through github (https://github.com/sanger-pathogens/snp-sites). The vcf file is available as supplementary Table S3.

Generalized linear model (GLM) construction (see equations 1 and 2), statistical analysis and tree visualizations was performed using R packages: dplyr, tidyr, ggplot2, ggtree, phytools, phangorn, lme4, dfoptim, car, reshape2, ggplot2, gridExtra, PredictABEL, dplyr, tidyr, scales, ggpubr.

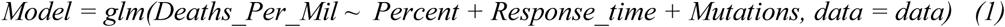

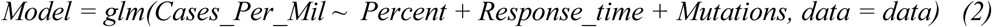

The *Percent* for each country *i* denotes the percentage occurrence for a specific mutation for that country and is calculated using the following formula:

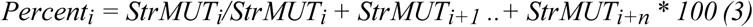

*Where: StrMUT* are the number of strains with the specific mutation for country *i* and *n* is the number of countries with at least one strain with the underlining mutation.

### Mutation tracking analyses

Relative frequencies across countries with at least one occurrence of the selected mutations of interest was visualized as bar plots across time (months). This provides an indication of the spread of the studied mutations across the general population. Analysis was performed using R (see packaged above).

### Additional Statistical analyses

Distributions of deaths per million for countries with the selected mutations and the reference mutation were visualized using box plots. Further statistical analysis was performed using standard non-parametric Wilcoxon test for comparing paired groups. The same analysis was performed for cases per million. Similarly, we performed the box plot visualizations for the same data but further separated by response time. We then applied the Kruskal–Wallis test for non-parametric method for testing whether the multiple groups originated from the same distribution. Analyses were performed using R (see packaged above).

### Structural prediction

Comparative structural modelling was carried out for unknown protein structures using the template-based web server, I-TASSER[28]. The accuracy of the method was assessed based on the I-TASSER confidence score (C-score), which indicates the quality of the predicted models. The C-score is calculated based on the significance of the sequence to structure alignments and the parameters of convergence obtained by the structure assembly simulations. It typically ranges between −5 and 2, and a value higher than −1.5 signifies a high accuracy of the predicted model[28]. I-TASSER was selected for protein structure modelling, since it outperformed other servers according to results from the 13th Community Wide Experiment on the Critical Assessment of Techniques for Protein Structure Prediction (CASP13)[30]. Protein structures were fixed, global energy (ΔG) was lowered when necessary, and mutagenesis was performed, using the FoldX4.0 suite (http://foldxsuite.crg.eu/). The PyMOL software (https://pymol.org/2/) was used for the visualization of the protein molecules.

RNA-protein docking simulations were carried out using the HDOCK[31] and MPRDock[32] algorithms. For docking simulations, as active and passive residues we selected residues described as such by Dinesh et al[27]. Protein-protein complexes were constructed using the ClusPro (v2.0)[33] and HDOCK[31] algorithms and binding affinities were calculated using the PRODIGY webserver[34]. Relative solvent accessibility (RSA) was calculated using the Missense3D[35] algorithm. Structural alignment was performed using the align tool of PyMOL and all-atom RMSD values were calculated without any outliers’ rejection, with zero cycles of refinement.

The DynaMut webserver[36], was used to visualize non-covalent molecular interactions, calculated by the Arpeggio algorithm[37]. SymmDock, a geometry-based docking algorithm for the prediction of cyclically symmetric complexes, was used for symmetric docking simulations of the REF-Q57H and ALT-Q57H homotetramers[38]. All docking simulations were performed in triplicates. Finally, binding affinities and dissociation constants (Kd) were calculated using the PRODIGY webserver[34].

## Notes

### Competing Interest Statement

The authors have declared no competing interest.

## References

1. Huang C, Wang Y, Li X, Ren L, Zhao J, Hu Y, et al. Clinical features of patients infected with 2019 novel coronavirus in Wuhan, China. Lancet. 2020. doi:10.1016/S0140-6736(20)30183-5

2. Gorbalenya AE, Baker SC, Baric RS, de Groot RJ, Drosten C, Gulyaeva AA, et al. The species Severe acute respiratory syndrome-related coronavirus: classifying 2019-nCoV and naming it SARS-CoV-2. Nature Microbiology. 2020. doi:10.1038/s41564-020-0695-z

3. Wu F, Zhao S, Yu B, Chen YM, Wang W, Song ZG, et al. A new coronavirus associated with human respiratory disease in China. Nature. 2020. doi:10.1038/s41586-020-2008-3

4. Andersen KG, Rambaut A, Lipkin WI, Holmes EC, Garry RF. The proximal origin of SARS-CoV-2. Nature Medicine. 2020. doi:10.1038/s41591-020-0820-9

5. Zhu N, Zhang D, Wang W, Li X, Yang B, Song J, et al. A novel coronavirus from patients with pneumonia in China, 2019. N Engl J Med. 2020. doi:10.1056/NEJMoa2001017

6. Lu R, Zhao X, Li J, Niu P, Yang B, Wu H, et al. Genomic characterisation and epidemiology of 2019 novel coronavirus: implications for virus origins and receptor binding. Lancet. 2020. doi:10.1016/S0140-6736(20)30251-8

7. Fehr AR, Channappanavar R, Perlman S. Middle East Respiratory Syndrome: Emergence of a Pathogenic Human Coronavirus. Annu Rev Med. 2017. doi:10.1146/annurev-med-051215-031152

8. Wu A, Peng Y, Huang B, Ding X, Wang X, Niu P, et al. Genome Composition and Divergence of the Novel Coronavirus (2019-nCoV) Originating in China. Cell Host Microbe. 2020. doi:10.1016/j.chom.2020.02.001

9. De Wit E, Van Doremalen N, Falzarano D, Munster VJ. SARS and MERS: Recent insights into emerging coronaviruses. Nature Reviews Microbiology. 2016. doi:10.1038/nrmicro.2016.81

10. Cui J, Li F, Shi ZL. Origin and evolution of pathogenic coronaviruses. Nature Reviews Microbiology. 2019. doi:10.1038/s41579-018-0118-9

11. Graham RL, Baric RS. Recombination, Reservoirs, and the Modular Spike: Mechanisms of Coronavirus Cross-Species Transmission. J Virol. 2010. doi:10.1128/jvi.01394-09

12. Petersen E, Koopmans M, Go U, Hamer DH, Petrosillo N, Castelli F, et al. Comparing SARS-CoV-2 with SARS-CoV and influenza pandemics. Lancet Infect Dis. 2020;3099: 1–7. doi:10.1016/S1473-3099(20)30484-9

13. Lv M, Luo X, Estill J, Liu Y, Ren M, Wang J, et al. Coronavirus disease (COVID-19): A scoping review. Eurosurveillance. 2020. doi:10.2807/1560-7917.ES.2020.25.15.2000125

14. Zhou P, Yang X Lou, Wang XG, Hu B, Zhang L, Zhang W, et al. A pneumonia outbreak associated with a new coronavirus of probable bat origin. Nature. 2020. doi:10.1038/s41586-020-2012-7

15. Khafaie MA, Rahim F. Cross-country comparison of case fatality rates of Covid-19/SARS-CoV-2. Osong Public Heal Res Perspect. 2020. doi:10.24171/j.phrp.2020.11.2.03

16. Chan-yeung M, Xu R. SARS: epidemiology CUMULATIVE NUMBER OF CASES AND DEATHS IN VARIOUS COUNTRIES IN. Respirology. 2003;8: S9–S14.

17. Petrosillo N, Viceconte G, Ergonul O, Ippolito G, Petersen E. COVID-19, SARS and MERS: are they closely related? Clinical Microbiology and Infection. 2020. doi:10.1016/j.cmi.2020.03.026

18. Wang C, Li W, Drabek D, Okba NMA, van Haperen R, Osterhaus ADME, et al. A human monoclonal antibody blocking SARS-CoV-2 infection. Nat Commun. 2020. doi:10.1038/s41467-020-16256-y

19. Katoh K. MAFFT: a novel method for rapid multiple sequence alignment based on fast Fourier transform. Nucleic Acids Res. 2002. doi:10.1093/nar/gkf436

20. Stamatakis A. RAxML version 8: A tool for phylogenetic analysis and post-analysis of large phylogenies. Bioinformatics. 2014. doi:10.1093/bioinformatics/btu033

21. Page AJ, Taylor B, Delaney AJ, Soares J, Seemann T, Keane JA, et al. SNP-sites: rapid efficient extraction of SNPs from multi-FASTA alignments. Microb genomics. 2016. doi:10.1099/mgen.0.000056

22. Korber B, Fischer W, Gnanakaran SG, Yoon H, Theiler J, Abfalterer W, et al. Spike mutation pipeline reveals the emergence of a more transmissible form of SARS-CoV-2. bioRxiv. 2020. doi:10.1101/2020.04.29.069054

23. Hale T, Webster, Sam, Anna Petherick, Toby Phillips BK. Oxford COVID-19 Government Response Tracker. In: Blavatnik School of Government. 2020.

24. Issa E, Merhi G, Panossian B, Salloum T, Tokajian S. SARS-CoV-2 and ORF3a: Nonsynonymous Mutations, Functional Domains, and Viral Pathogenesis. mSystems. 2020. doi:10.1128/msystems.00266-20

25. Pachetti M, Marini B, Benedetti F, Giudici F, Mauro E, Storici P, et al. Emerging SARS-CoV-2 mutation hot spots include a novel RNA-dependent-RNA polymerase variant. J Transl Med. 2020. doi:10.1186/s12967-020-02344-6

26. Hadfield J, Megill C, Bell SM, Huddleston J, Potter B, Callender C, et al. NextStrain: Real-time tracking of pathogen evolution. Bioinformatics. 2018. doi:10.1093/bioinformatics/bty407

27. Dinesh DC, Chalupska D, Silhan J, Veverka V, Boura E. Structural basis of RNA recognition by the SARS-CoV-2 nucleocapsid phosphoprotein. bioRxiv. 2020; 2020.04.02.022194. doi:10.1101/2020.04.02.022194

28. Zhang Y. I-TASSER server for protein 3D structure prediction. BMC Bioinformatics. 2008. doi:10.1186/1471-2105-9-40

29. Kang S, Yang M, Hong Z, Zhang L, Huang Z, Chen X, et al. Crystal structure of SARS-CoV-2 nucleocapsid protein RNA binding domain reveals potential unique drug targeting sites. Acta Pharm Sin B. 2020. doi:10.1016/j.apsb.2020.04.009

30. Guzenko D, Lafita A, Monastyrskyy B, Kryshtafovych A, Duarte JM. Assessment of protein assembly prediction in CASP13. Proteins Struct Funct Bioinforma. 2019. doi:10.1002/prot.25795

31. Yan Y, Tao H, He J, Huang SY. The HDOCK server for integrated protein–protein docking. Nat Protoc. 2020. doi:10.1038/s41596-020-0312-x

32. He J, Tao H, Huang SY. Protein-ensemble-RNA docking by efficient consideration of protein flexibility through homology models. Bioinformatics. 2019. doi:10.1093/bioinformatics/btz388

33. Vajda S, Yueh C, Beglov D, Bohnuud T, Mottarella SE, Xia B, et al. New additions to the ClusPro server motivated by CAPRI. Proteins Struct Funct Bioinforma. 2017. doi:10.1002/prot.25219

34. Xue LC, Rodrigues JP, Kastritis PL, Bonvin AM, Vangone A. PRODIGY: A web server for predicting the binding affinity of protein-protein complexes. Bioinformatics. 2016. doi:10.1093/bioinformatics/btw514

35. Ittisoponpisan S, Islam SA, Khanna T, Alhuzimi E, David A, Sternberg MJE. Can Predicted Protein 3D Structures Provide Reliable Insights into whether Missense Variants Are Disease Associated? J Mol Biol. 2019. doi:10.1016/j.jmb.2019.04.009

36. Rodrigues CH, Pires DE, Ascher DB, RenéRen I, Rachou R, Oswaldo Cruz F. DynaMut: predicting the impact of mutations on protein conformation, flexibility and stability. Nucleic Acids Res. 2018;46. doi:10.1093/nar/gky300

37. Jubb HC, Higueruelo AP, Ochoa-Montaño B, Pitt WR, Ascher DB, Blundell TL. Arpeggio: A Web Server for Calculating and Visualising Interatomic Interactions in Protein Structures. J Mol Biol. 2017;429: 365–371. doi:10.1016/j.jmb.2016.12.004

38. Schneidman-Duhovny D, Inbar Y, Nussinov R, Wolfson HJ. PatchDock and SymmDock: servers for rigid and symmetric docking. doi:10.1093/nar/gki481

