## Supplementary Figures for "Generalized linear models provide a measure of virulence for specific mutations in SARS-CoV-2 strains"

**A**

**B**

**C**

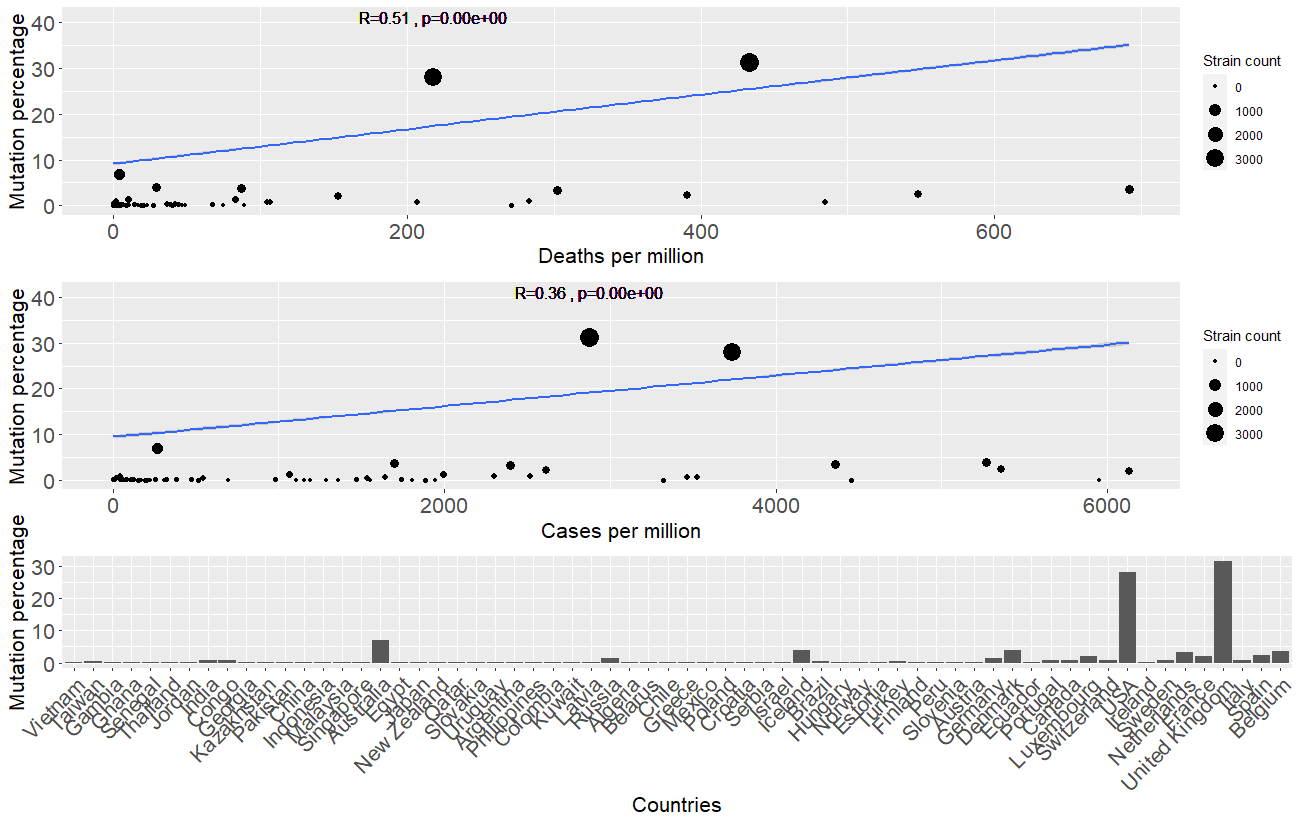

**Fig S1**. Analyses plots for S protein mutation at position 23403 (D614G) **A**. Regression model line showing the simplified fit for mutations percentage across countries and the death rate per million for each country. Pearson’s correlation is shown by the *R* value accompanied by the *p*-value of the correlation coefficient. **B**. Similar regression fit for mutations percentage across countries this time showing cases per million for each country. **C**. Detailed histogram of the percentage occurrence of the mutation across different countries. Countries are sorted with increasing deaths per million.

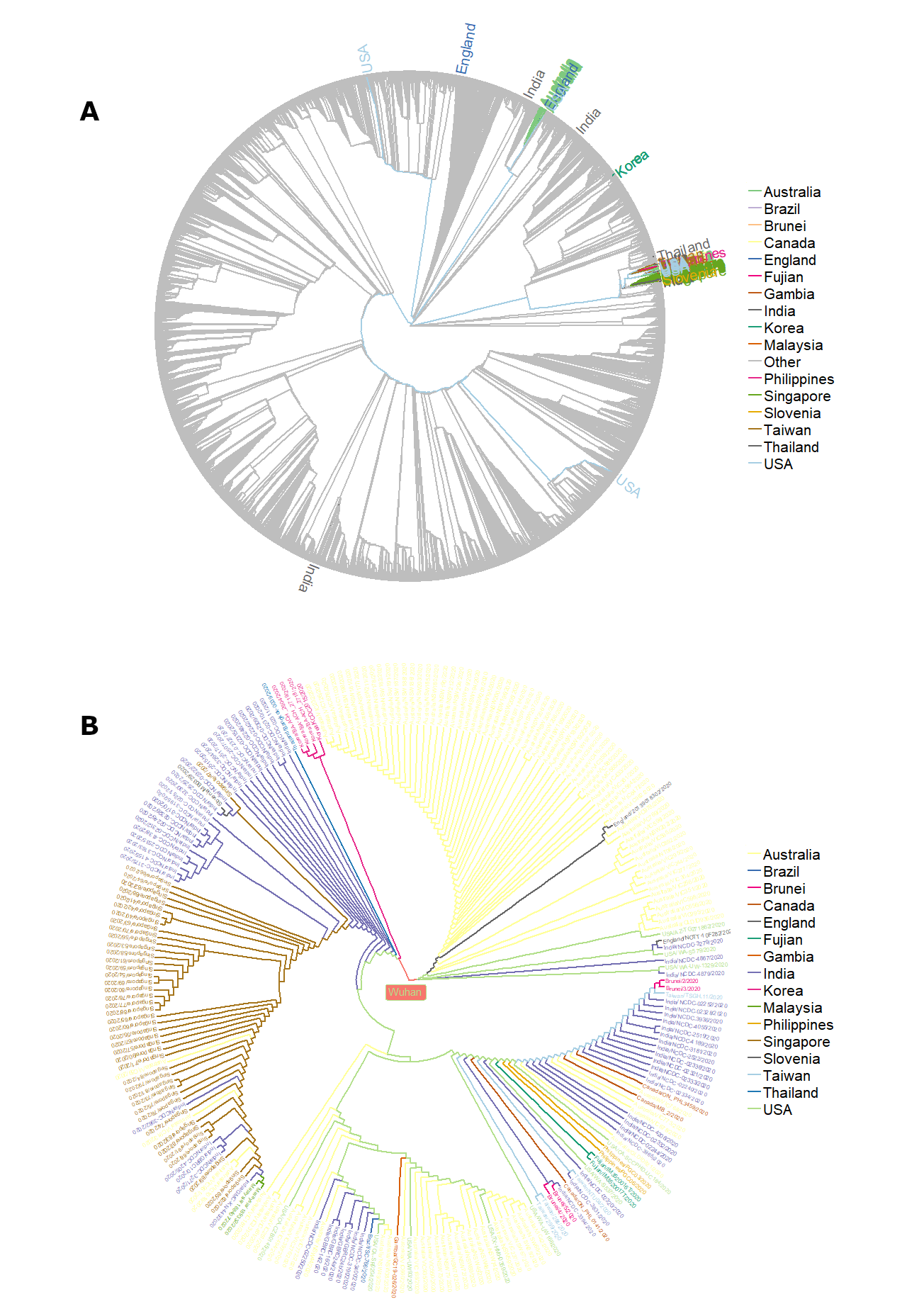

**Fig S2. Phylogenetic Trees using 16,535 trimmed full genome SARS- CoV-2 strains from GISAID**. **A.** A maximum likelihood (RAxML) tree, exapandig form the Wuhan reference strain, with the strains comprising the P13L clade highlighted in colour. Branches of the tree are colour coded by country. **B.** Close up of the P13L clade expanded to show patterns of the

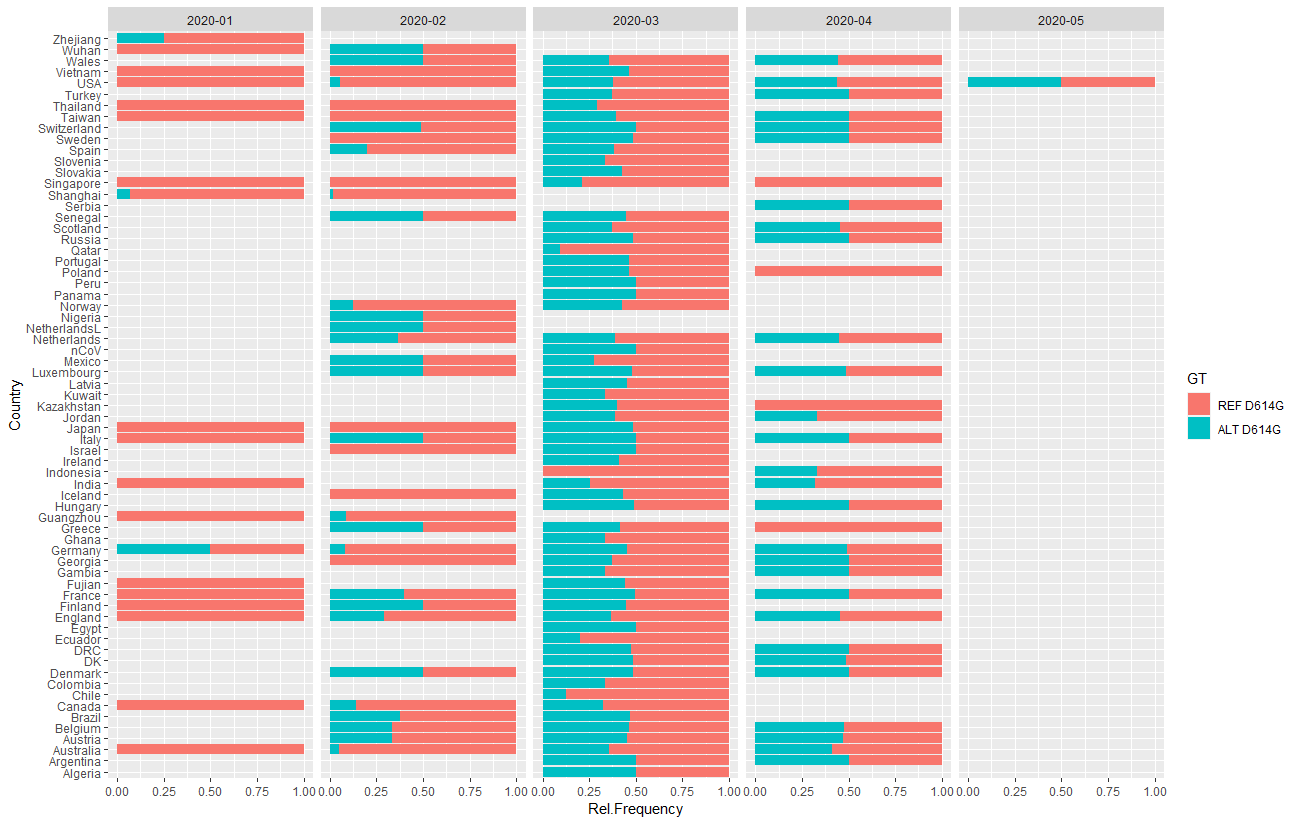

**Fig S3.** D614G transmission in months from its first occurrence in Europe (Germany) to its transmission across the globe.

P13L mutation that is being tracked across the phylogenetic tree. **C.** Same RAxML tree as in A, with the strains comprising the Q57H clade highlighted in colour. Branches of the tree are colour coded by country.

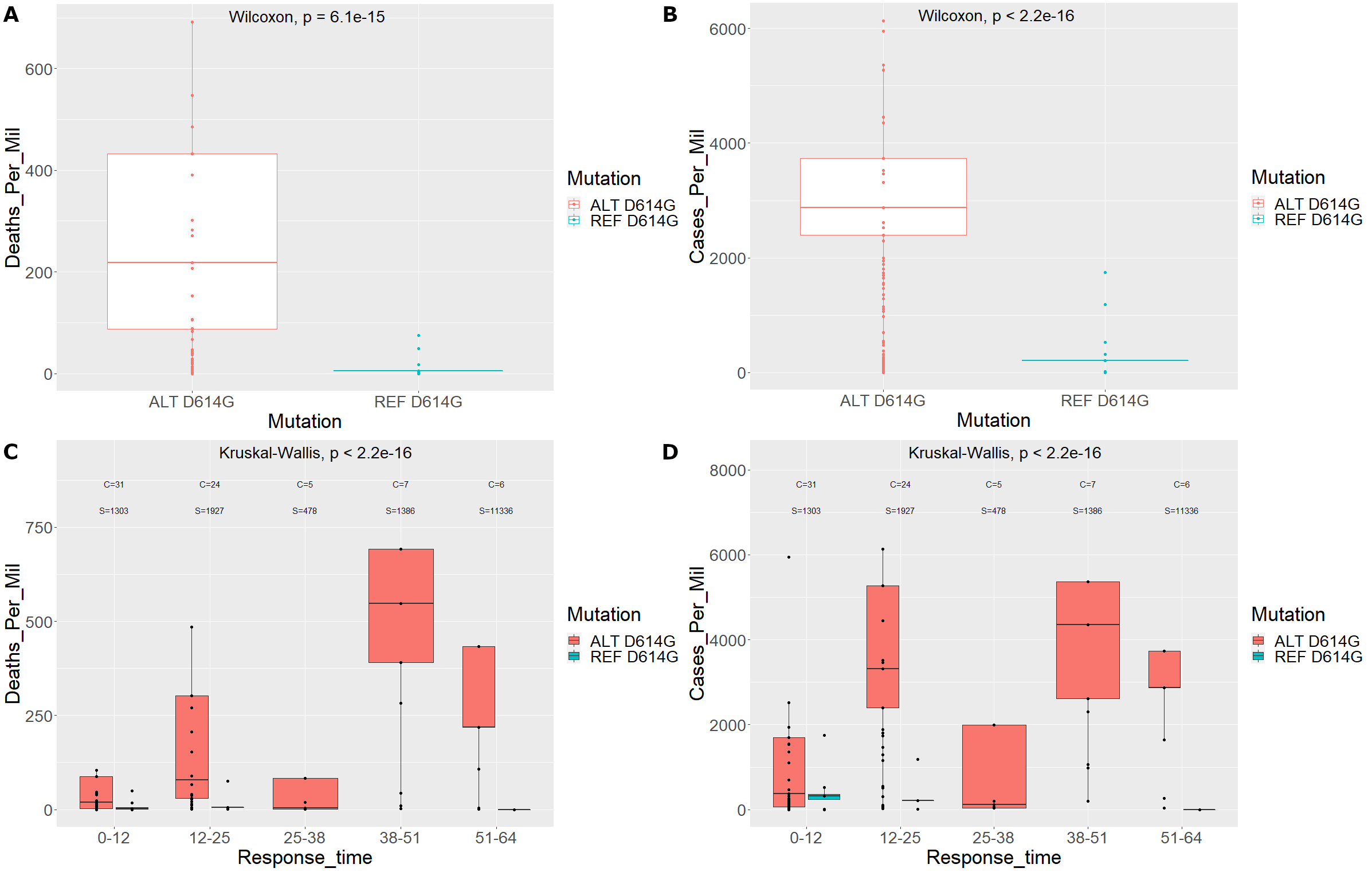

**Fig S4**. Boxplot distributions with and without the D614G mutation. **A.** Deaths per million for countries with the D614G mutation and the reference mutation. **B.** Cases per million for countries with the D614G mutation and the reference mutation. **C.** Deaths per million for countries with the D614G mutation and the reference mutation including response time separation. *C* denotes the number of unique countries in the group and *S* is the number of strains in the group. **D.** Cases per million for countries with the D614G mutation and the reference mutation including response time separation. *C* and *S* are as denoted for panel **C**.
